# End-To-End, Fully Automated Manufacturing of Hematopoietic Stem and Progenitor Cell Therapy Via*CCR5*-Targeted Integration

**DOI:** 10.64898/2026.10.28.758978

**Authors:** Jack M.P. Castelli, Nowah Afangbedji, Sofia E. Luna, Freja K. Ekman, Dmitry Malkov, Karthik Shastry, Victor Rocha, Arpit Batish, Anna Sahakyan, Julie Fang, Zhou Li, Kayla Ruggiero, Jessica Fioravanti, Fabrizia Urbinati, Matthew H. Porteus

## Abstract

Gene therapies utilizing hematopoietic stem and progenitor cells (HSPCs) have curative potential across hematologic diseases, but manual manufacturing workflows are costly and difficult to scale, limiting patient access. Here, we demonstrated automation of an HSPC gene knock-in protocol on a high-throughput manufacturing platform designed for clinical-scale production. The platform executed all steps, from post-thaw HSPC inoculation through Cas9 electroporation, AAV6 transduction, and harvest, producing gene-edited HSPCs (geHSPCs) carrying an anti-HIV antibody transgene at the *CCR5* locus with high purity, preserved differentiation potential, and editing efficiencies comparable to small-scale processing. Following transplantation into mice, geHSPCs engrafted and reconstituted multiple hematopoietic lineages, demonstrating that repopulation capacity was preserved. This work shows that clinical-scale doses of geHSPCs meeting release criteria can be produced on an automated, high-throughput platform, while preserving key product attributes and *in vivo* function, providing a path toward scalable and cost-effective manufacturing that could broaden access to HSPC gene therapies.

**GRAPHICAL ABSTRACT:** 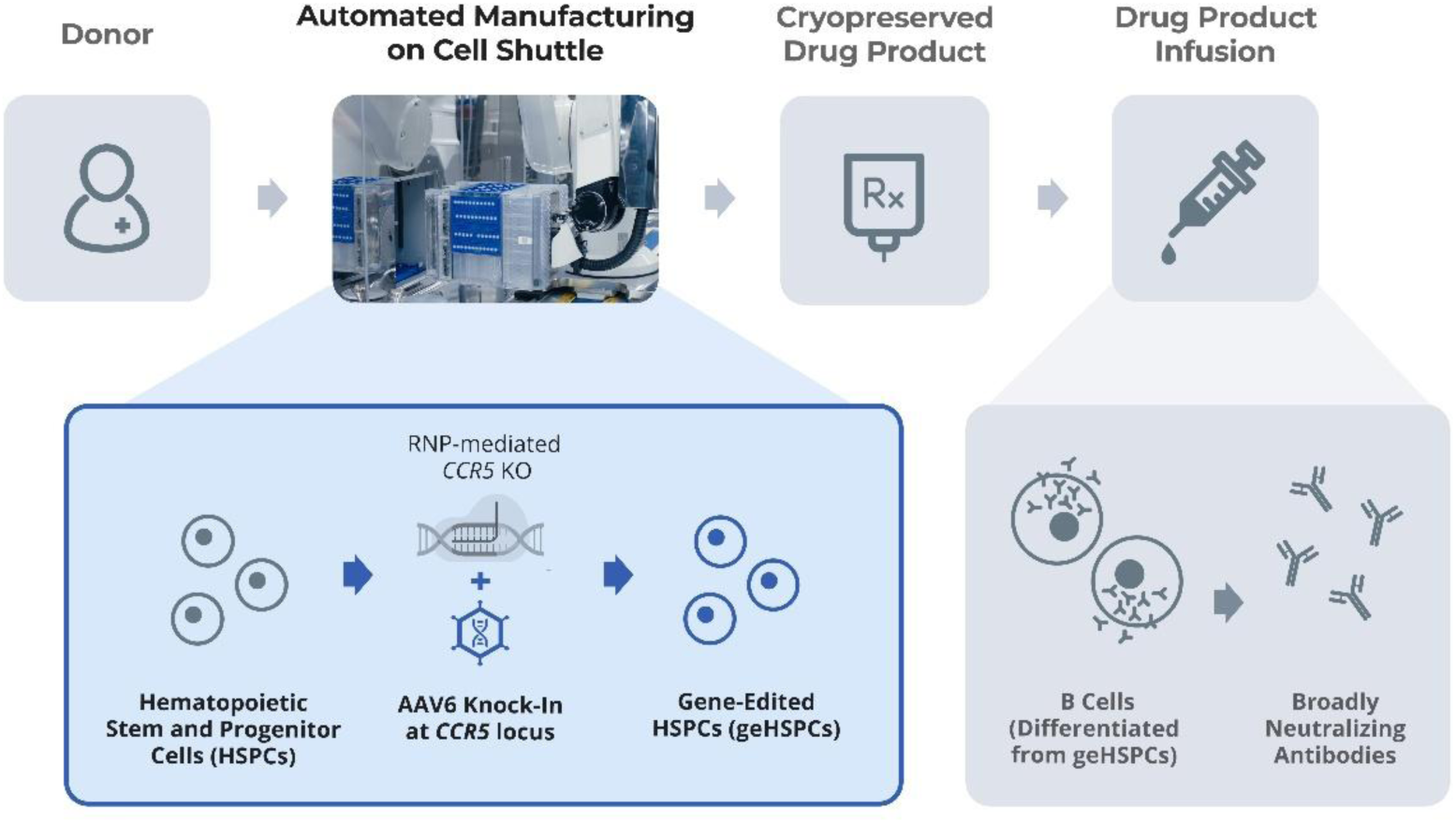

## Introduction

Transplantation of hematopoietic stem and progenitor cells (HSPC) treated with gene therapies has become a clinically validated route offering durable, potentially curative treatments for hematologic and immunologic disorders, with seven approved products and more than 50 active clinical trials.^1^ An expanding fraction of these autologous therapies are based on genome editing of HSPCs (geHSPCs). Therapeutic transgenes can be introduced by CRISPR-Cas targeted knock-in (KI), achieved through homology-directed repair (HDR) in the presence of a repair DNA template.^2^ HIV is one setting in which this approach is being pursued, where transgene KI in HSPCs yields broadly neutralizing antibody (bnAb) expression in B cell progeny at levels sufficient to neutralize HIV.^3,4^ Targeting the *CCR5* locus for KI simultaneously disrupts the major HIV-1 co-receptor through knockout (KO) events, adding a layer of resistance to HIV infection in the transplanted cells and their progeny.^5^

The cost and complexity of manufacturing geHSPC products remain a central barrier to the accessibility of these therapies. Approved treatments for sickle cell disease and transfusion-dependent β-thalassemia cost millions of dollars per patient in the US and require large CD34^+^ cell collections obtained over multiple apheresis cycles to offset variable manufacturing yields.^6,7^ Despite the continued clinical success of these programs, patients enrolled in trials wait for a median of four months or longer between cell collection and reinfusion, and some discontinue treatment altogether because of manufacturing failures.^8–11^ Partially automated manufacturing at Good Manufacturing Practice (GMP) facilities reduces hands-on processing time and batch-to-batch variability but does not remove the fixed and marginal costs that constrain scale.^12,13^

Centralized manufacturing with fully automated and closed workflows, such as those widely adopted for other classes of biologics, offers a route to reducing total costs with high-throughput, parallel processing.^14^ Lowering the per-dose cost is a prerequisite for meeting market demands and expanding access, particularly for indications such as HIV, where the largest burden of disease falls on low-and middle-income countries, in which point-of-care approaches are prohibitively expensive and infrastructure-intensive.^15,16^ To this end, a 5-day process for producing geHSPCs was developed on the Cell Shuttle^®^ (Cellares), an automated, high-throughput manufacturing platform. Its closed, single-use consumable cartridge contains integrated components required for geHSPC manufacturing, including bioreactors for cell culture and electroporation (EP) chambers for delivery of editing reagents. Here, we show that clinical-scale doses of geHSPCs can be manufactured while preserving editing levels and key HSPC properties, and we estimate that this approach significantly reduces the cost of the geHSPC product. This work establishes a scalable manufacturing path that reduces manual intervention, cost, and infrastructure burdens that currently limit access to HSPC gene therapies.

## RESULTS

### Engineering process for gene KI using an automated manufacturing platform

The Cell Shuttle has been deployed for CAR T cell manufacturing in ongoing clinical trials, but its suitability for HSPCs has not been established. Our first objective was to define culture conditions for CD34^+^ HSPCs within the single-use consumable cartridge (CC). Adapting the research-scale process to the platform required no substantive changes. Cryopreserved CD34^+^ hematopoietic stem and progenitor cells (HSPCs) were cultured for three days prior to EP and recovered for two days prior to formulation, as previously described.^5^ Manual intervention was limited to initial cell thawing and transfer onto the single-use CC, and formulation of critical reagents, with all subsequent manufacturing steps performed on the Cell Shuttle (Figure 1).

**Figure 1.**
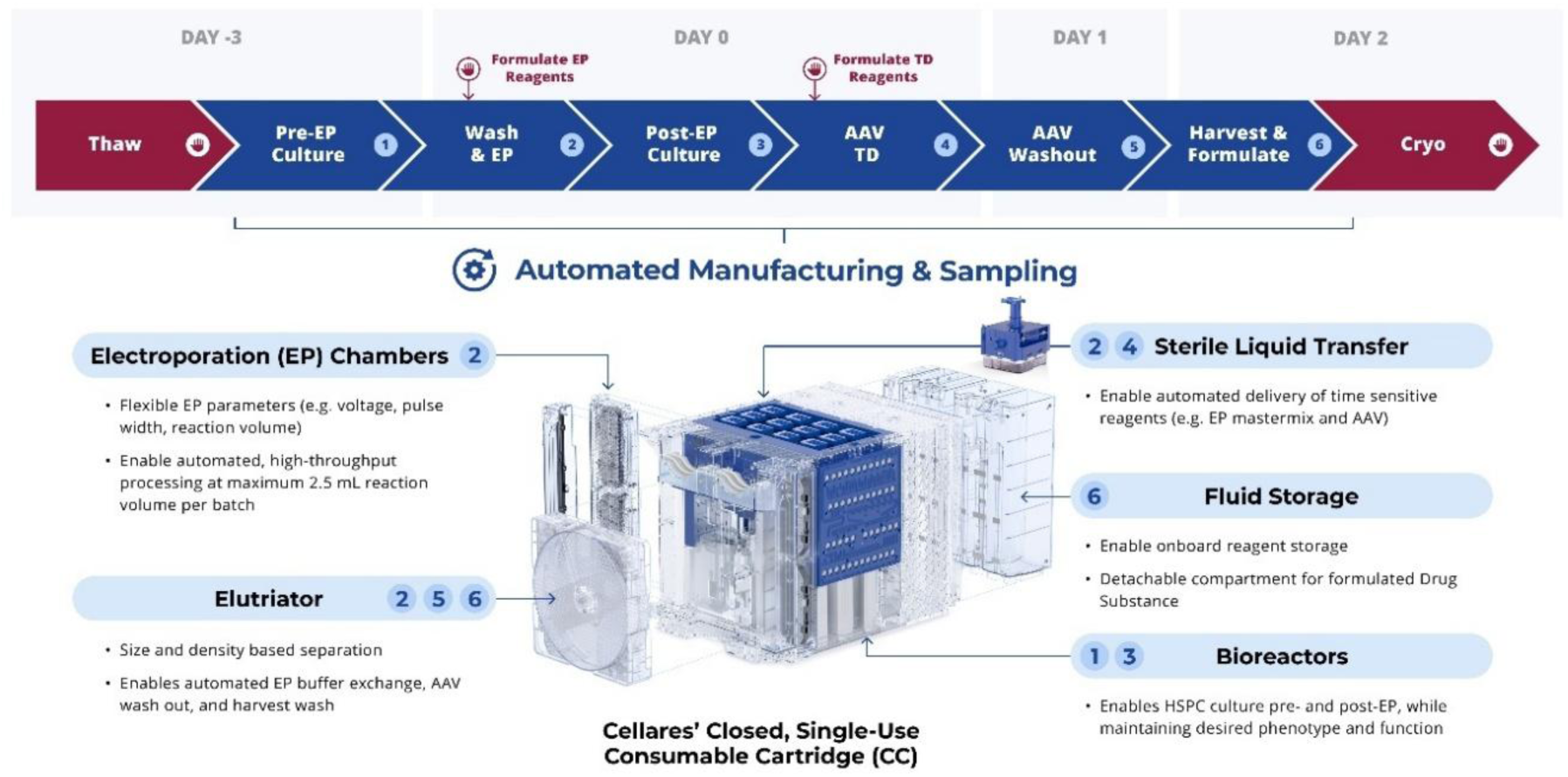
Overview of the automated process for engineering CD34^+^ cells. Cryopreserved CD34⁺ HSPCs were thawed on day −3 and cultured for 3 days under pre-electroporation (Pre-EP) conditions, followed by EP on Day 0. Cells were subsequently cultured under post-EP conditions in the presence of AAV, followed by AAV washout on day 1, continued culture, and final harvest, formulation, and cryopreservation on day 2. Numbered steps correspond to the following conditions: (1) Pre-EP culture: 37°C, 5% CO₂, 5% O₂ (hypoxia), static culture, seeding density 3 × 10⁶ cells/cm² (0.6 × 10⁶ cells/mL); (2) EP: 450 V, 400 μs, 4 s, 2 pulses; (3) Post-EP culture: 37°C, 5% CO₂, 5% O₂ (hypoxia), static culture, seeding density 10 × 10⁶ cells/cm² (1 × 10⁶ cells/mL); (4) AAV6 transduction: MOI 1,250 for 24 h; (5, 6) Harvest and formulation by counterflow centrifugal elutriation (CCE). All steps are carried out within Cellares’ closed, single-use Consumable Cartridge (CC). Callouts indicate the cartridge compartment for each numbered operation: Bioreactors (steps 1, 3), EP Chambers (step 2), Elutriator (steps 2, 5, 6), Sterile Liquid Transfer (steps 2, 4) and Fluid Storage (step 6). Steps requiring operator intervention are indicated by the hand icon, fully automated steps are indicated by the gear icon. Arrows indicate key manual interventions, including reagent preparation and HSPC thawing, RNP formulation prior to EP, AAV6 preparation prior to transduction, automated sampling during harvest, and DMSO addition during fill-finish prior to cryopreservation.

To edit HSPCs, we knocked in an anti-HIV bnAb transgene (10-1074) at the *CCR5* locus using Cas9 ribonucleoprotein (RNP), with the single-guide RNA (sgRNA) and 4.2 kb AAV6 repair template unchanged from the research-scale process (Figure 2A).^17^ The research-scale process was established at 1.0×10^7^ live cells per mL in 100 µL reaction volume. To achieve our clinical manufacturing target of at least 3.0×10^8^ HSPCs electroporated per manufacturing run, we therefore needed to optimize parameters on the Cellares EP platform, including increasing the cell concentration (Figure 2B).

**Figure 2.**
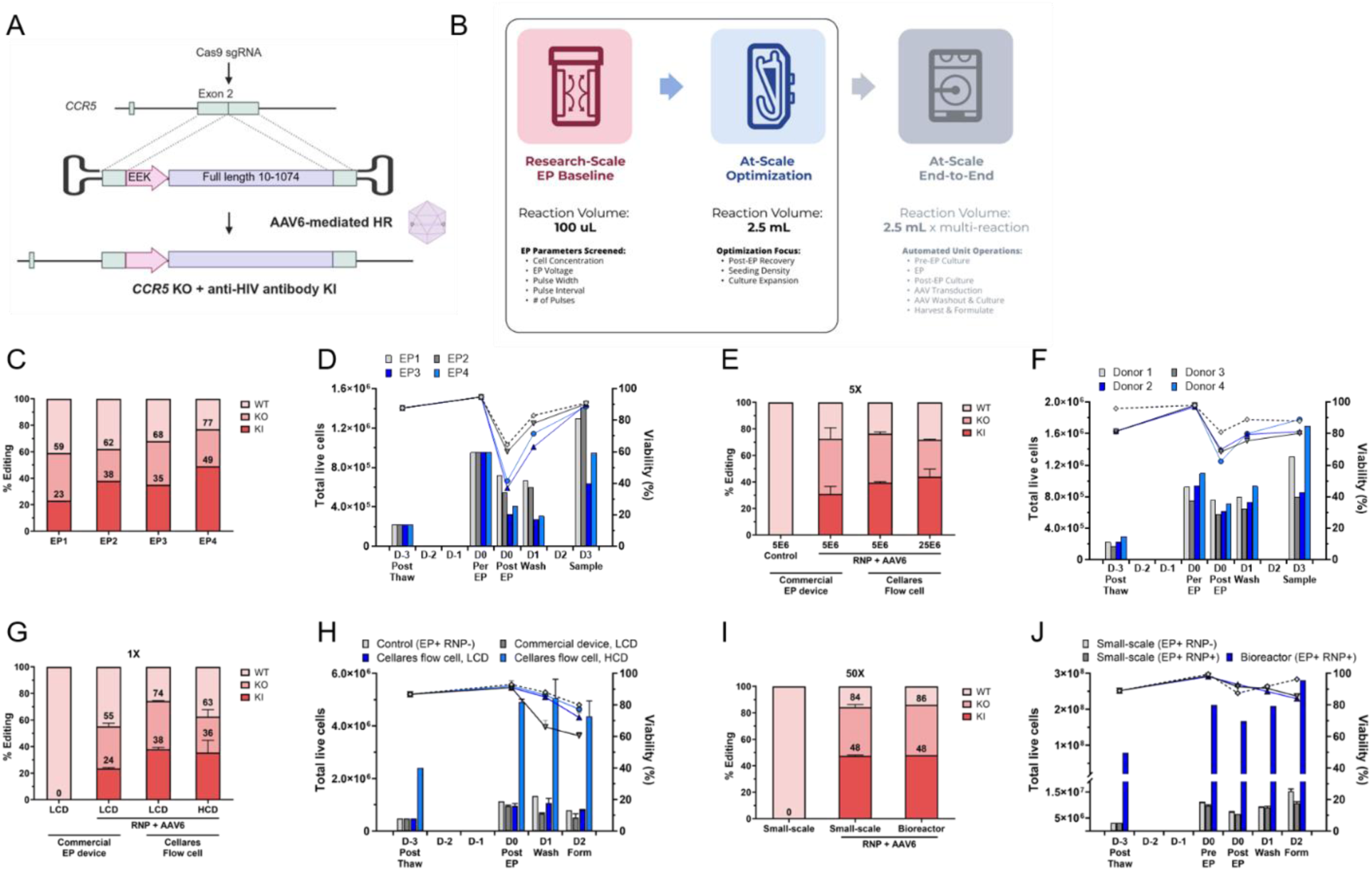
Automated platform supports growth and editing of HSPCs at high density. **(A)** Schematic of the gene-editing strategy targeting the *CCR5* locus. **(B)** Overview of the EP process scale-up strategy from Research Baseline EP-scale to At-Scale Optimization; At-Scale End-to-End is presented in Fig. 3. (C, D) EP parameter screening in human HSPCs using the Cellares EP System across four conditions (1×10⁷ cells/mL, 100uL reactions) differing in voltage, pulse width, pulse interval, and number of pulses. (C) Gene-editing efficiency, including knockout (KO) and knock-in (KI). (D) Total live cells (bars, left axis) and viability (lines, right axis) from day-3 post-thaw through day 2. (E, F) Evaluation of the optimized EP4 condition across four independent healthy donors. (E) KO and KI efficiencies. (F) Total live cells and viability across the same process time points shown in (D). (G, H) Editing performance and cell health at small EP scale (1x: 100 µL), comparing low cell density (LCD: 1×10⁷ cells/mL) and high cell density (HCD: 5×10⁷ cells/mL) conditions. (G) KO and KI efficiencies comparing a commercial EP device with the Cellares flow cell with or without RNP and AAV6. (H) Total live cells per reaction and viability across the same process time points shown in (D). (I, J) Editing performance and cell health following scale-up to the bioreactor EP-scale (50x: 5 mL). (I) KO and KI efficiencies comparing small-scale, bioreactor and EP only control. (J) Total live cells and viability. Data are presented as mean ± standard deviation (SD); n = 2 technical replicates from one donor.

We first screened EP voltage, pulse width, pulse interval, and number of pulses at the research-scale cell concentration to identify the optimal EP condition (Figure 2C-D). EP4 achieved the highest total editing efficiency (49% KI and 77% total KO, Figure 2C), while maintaining viability and expansion comparable to the other EP conditions (Figure 2D). Therefore, EP4 was selected for subsequent studies. EP4 showed reproducible performance across four independent healthy donors, with 69–79% total editing (Figure 2E) and comparable cell viability and expansion kinetics (Figure 2F), supporting the reproducibility of the optimized EP condition.

We next translated this optimized condition to higher cell concentration to enable scale-up. *CCR5* editing and cell expansion were within expected ranges: Template KI on the Cellares electroporator system (EPS) reached 38% and 36% of alleles at low EP cell density (LCD, 1.0×10^7^ live cells per mL) and high EP cell density (HCD, 2.5×10⁶ live cells per mL), respectively, compared with 24% KI on a commercial EP device. Total KO ranged from 63% to 74% of alleles on the Cellares EP System, compared with 55% on the commercial device (Figure 2G). Post-EP cell growth and viability were comparable to control HSPCs electroporated without RNP and without AAV6 transduction: 1.66-fold and 1.81-fold expansion between day –3 (post-thaw) and day 2 (formulation) for LCD and HCD, respectively, versus 1.64-fold for the control, with viability ≥70% (≥80% for the control). EP with the commercial device reduced expansion to 1.07-fold and viability to 60% (Figure 2H).

We then evaluated the optimized condition at a clinically relevant cell concentration of 5.0×10⁷ cells per mL at 2.5 mL reaction volume per batch (50×). This condition was initially evaluated using the EP unit operation alone, with only two reactions performed at a time to minimize cell and reagent consumption (Figure 2I-J, Figure S1A-B). Editing and cell expansion remained comparable to control conditions. Editing levels were similar to those at smaller scales, at 48% KI and 84%–86% KO (Figure 2I). Expansion at small-scale in the Cellares flow cell was 3.38-fold, 3.57-fold in the bioreactor, and 4.86-fold for the control (≥80% viability; Figure 2J). Because larger numbers of HSPCs could be cultured at HCD without loss of editing, purity, or expansion, subsequent manufacturing runs on the CC were performed at HCD (Figure S1C-E).

### Production of clinical doses using mobilized CD34^+^ cells from healthy donors

To demonstrate clinical-scale manufacturing, CD34^+^ cells were collected from the mobilized peripheral blood (mPB) of healthy adult donors and used in two automated manufacturing runs, each targeting a minimum of 3.0×10^8^ HSPCs electroporated per run. This required 5 consecutive 2.5 mL EP reactions at 5.0×10^7^ HSPCs/mL (Figure 3A).

**Figure 3.**
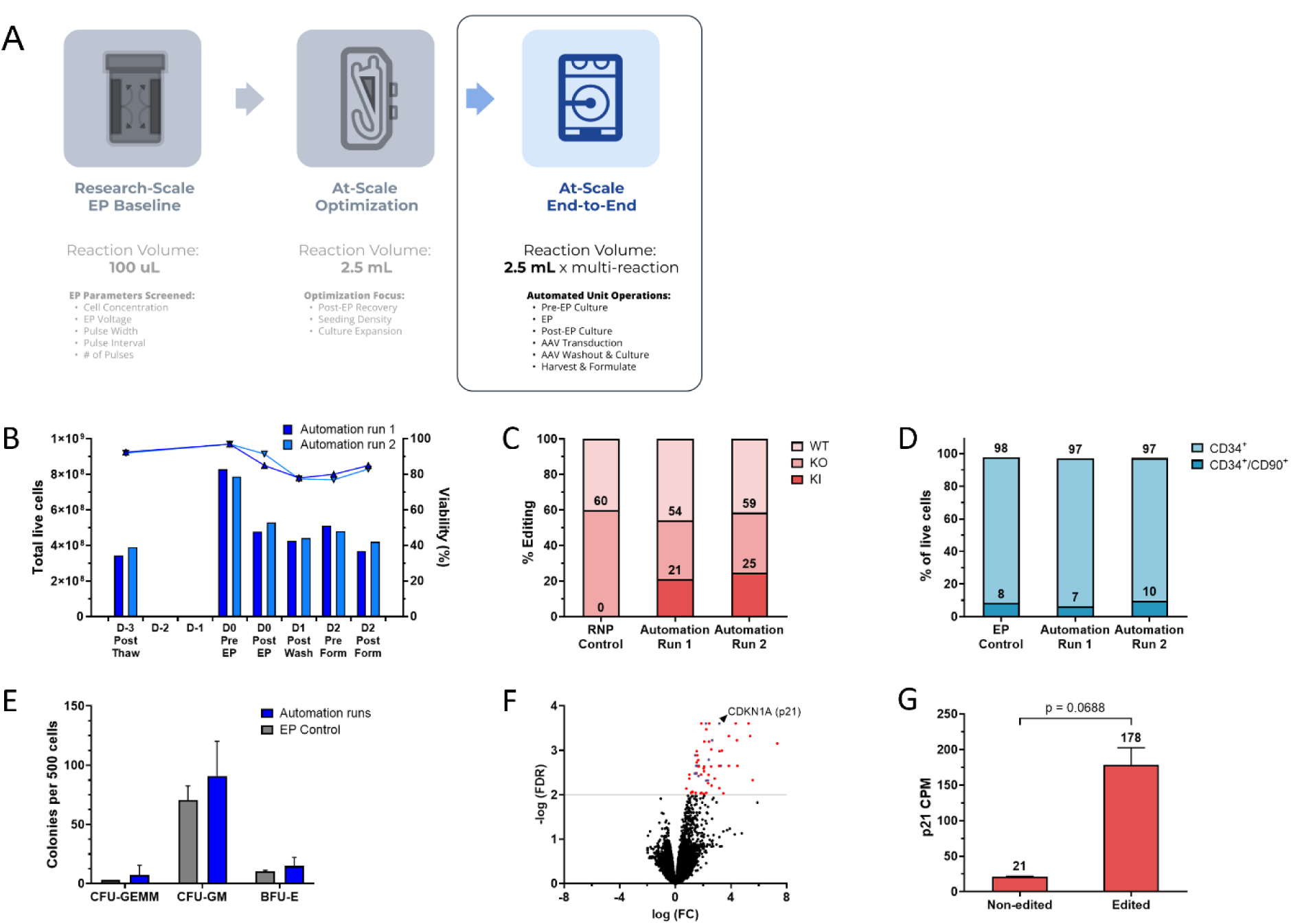
Clinical dose of HSPCs edited at high efficiency while retaining stemness and differentiation potential. **(A)** Process scale-up schematic highlighting (5) Clinical Scale Benchtop and (6) Clinical Scale in the Automated Cellares Cell Shuttle® (E2E on Consumable Cartridge), continuing from the scale-up strategy shown in Fig. 2. **(B)** Total live cell counts and viability across the process (day −3 post-thaw through day 2 post-formulation) for two independent automation runs. **(C)** Gene-editing outcomes (% WT, KO, KI) at the *CCR5* locus for RNP-only control versus two automated Cell Shuttle runs. **(D)** Frequency of CD34^+^ and CD34⁺/CD90⁺ cells by flow cytometry for EP control and automation runs. **(E)** Number of colonies formed per 500 cells plated in colony forming unit (CFU) assays for EP control and automation runs. CFU-GEMM: CFU-granulocyte, erythroid, macrophage, megakaryocyte. CFU-GM: CFU granulocyte and monocyte. BFU-E: burst forming unit-erythroid. Mean and SD plotted; n = 2 technical replicates for EP control; n = 3 technical replicates for automation runs. No statistically significant differences by two-way ANOVA (p > 0.05). **(F)** Volcano plot depicting differentially expressed genes between non-edited (EP and RNP controls) and edited (automation runs) HSPCs as measured by fold-change (FC) from RNA-seq. Red dots show genes with significant false discovery rates (FDR < 0.01), with purple dots showing genes belonging to the P53 response pathway. **(G)** Normalized p21 (*CDKN1A*) expression as measured by counts per million (CPM). Mean and SD are plotted; n = 2 replicates; significance calculated by unpaired two-tailed *t* test.

Both runs were completed without significant deviation from protocol and met target release criteria for cell count, viability, and CD34^+^ purity (Table 1). Starting from 3.43×10⁸ and 3.89×10⁸ live cells, the runs produced 107%–108% final yield with 83.0%–85.0% viability at formulation (Figure 3B). Absolute yields of 3.68×10⁸ and 4.21×10⁸ live cells were harvested and formulated as final drug product, sufficient to each dose a 70 kg adult at over 5×10⁶ cells per kg, a relevant clinical threshold.^1^ Cryopreserved geHSPCs were shipped from the Cellares manufacturing facility to Stanford, analogous to clinical autologous cell products for bedside infusion. Post-thaw, *CCR5* editing reached 21.1%–24.7% KI and 54.2%–58.6% KO (Figure 3C). KI levels were comparable to those observed at research-scale at the same low AAV6 doses and without HDR enhancers, and total KO levels were comparable to a control EP reaction using RNP only.

**Table 1.** Target release criteria.

|  | Target | Run 1 | Run 2 |
| --- | --- | --- | --- |
| Viability (% live cells) | ≥70% | 75% | 73% |
| Purity (% CD34<sup>+</sup>) | ≥80% | 90.5% | 87.5% |
| CCR5 Editing (% KO) | ≥50% | 54.2% | 58.6% |
| CCR5 Editing (% KI) | ≥20% | 21.1% | 24.7% |
| Cell Dose (viable cells/kg)<br>Setpoint: 70kg | ≥5×10<sup>6</sup> | 5.26×10<sup>6</sup> | 6.01×10<sup>6</sup> |

### Phenotype of geHSPCs following automated manufacturing process

To characterize geHSPCs in the final drug product, we assessed cellular phenotype and HSPC function. We measured CD34^+^ and CD34^+^/CD90^+^ purity to identify the HSPC subsets responsible for hematopoietic reconstitution and multilineage engraftment.^18^ CD34^+^ purity remained high after manufacturing, at 97.0% and 97.3% for the two automation runs, versus 97.8% for the EP control. The CD34^+^/CD90^+^ populations were within typical ranges at 6.5% and 9.8%, versus 8.3% for the EP control (Figure 3D). We next assessed the stemness and differentiation potential of geHSPCs. Colony forming unit (CFU) assays showed no significant difference in the total number or distribution of colonies between the automation runs and the EP control (Figure 3E).

To evaluate toxicity at the transcriptional level, we performed bulk RNA sequencing on geHSPCs immediately post-thaw, given that Cas9/AAV6-mediated HDR has been associated with activation of DNA damage response pathways that could impair HSPC engraftment. Sixty genes were significantly upregulated in edited samples from the automation runs relative to unedited samples. Thirteen of these (21.7%) were associated with the p53 response pathway, with *CDKN1A* and *MDM2* showing the largest changes (Figure 3F). *CDKN1A* encodes p21, a transcriptional target of p53 with a role in HSPC cell cycle regulation; p21 abundance was upregulated 8.48-fold, though additional runs would be required to establish true significance (Figure 3G). GO term enrichment analysis revealed seven downregulated pathways and 22 upregulated pathways, with a number overlapping with the p53 response pathway, such as inflammatory response and apoptosis (Table S1).

These data describe the *in vitro* phenotype of geHSPCs following automated manufacturing at clinical scale. To evaluate the persistence of stem-like features and editing levels *in vivo*, we next performed engraftment studies.

### Transplantation of final drug product into immunodeficient mice

Clinical translation to hematopoietic stem cell transplantation (HSPCT) requires that CD34^+^ cells retain engraftment capacity throughout manufacturing. We therefore transplanted geHSPCs into NBSGW mice and followed hematopoietic reconstitution *in vivo*. Adult NBSGW mice support engraftment of human CD34^+^ cells with low-dose busulfan conditioning.^19,20^ We transplanted a limited number of cells (2.5×10⁵ per mouse) to resolve any difference in engraftment between edited and unedited HSPCs (Figure 4A). Engraftment of multiple hematopoietic lineages and editing levels were tracked in peripheral blood (PB) starting at 8 weeks post-transplantation and across tissues at necropsy 12 weeks post-transplantation.

**Figure 4.**
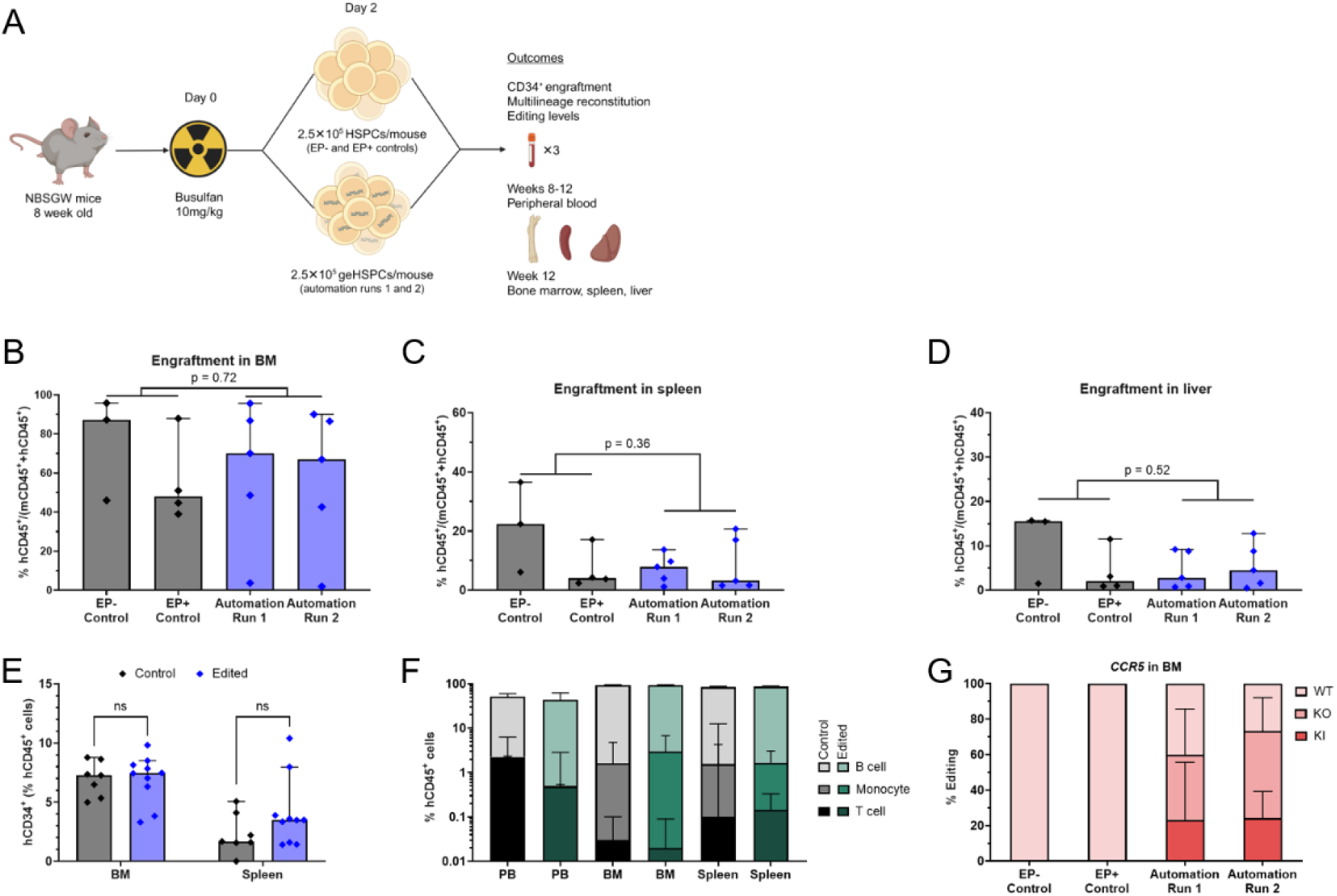
Engraftment and multi-lineage reconstitution preserved in edited HSPCs. **(A)** Experimental outline of engraftment study in busulfan-conditioned, adult NBSGW mice. 24 hours post-thaw, 2.5 × 10^5^ HSPCs were injected intravenously per mouse. Starting at week 8, peripheral blood (PB) was collected every 2 weeks and, at week 12 post-transplantation, tissues were recovered for analysis. **(B-D)** Engraftment in bone marrow (BM), spleen, and liver between control HSPCs (EP-and EP+) and geHSPCs (automation runs) as measured by flow cytometry on human and mouse CD45^+^ cells. n = 3-5 mice per condition. **(E)** HSPC populations in BM and spleen as measured by flow cytometry on CD34^+^ cells. n = 7-10 mice per group. **(F)** Hematopoietic lineages in PB, BM, and spleen as measured by flow cytometry on CD19^+^/CD3^-^cells (B cell), CD3^+^/CD19^-^cells (T cell), and CD33^+^/CD3^-^/CD19^-^cells (monocytes). n = 7-10 mice per group. No statistically significant differences by two-way ANOVA (p > 0.05). **(G)** Gene-editing outcomes at *CCR5* locus as measured by sequencing genomic DNA from BM samples. n = 3-5 mice per condition. Median and 95% confidence interval (CI) are plotted in B-F; mean and SD are plotted in G; significance calculated by unpaired two-tailed *t* test in B-E.

Circulating human CD45^+^ (hCD45^+^) cells in PB were modest between weeks 8 and 12 (0.180%–7.83% of total mouse and human CD45^+^ white blood cells) (Figure S2A). At necropsy, hCD45+ populations were detected in bone marrow (BM), spleen, and liver. BM engraftment reached a median of 67.0% and 70.1% for the two automation runs, versus 87.2% and 47.9% for the non-electroporated (EP-) and electroporated (EP+) controls, respectively (Figure 4B). Engraftment was likewise observed in the spleen (3.20% and 7.90% versus 22.4% and 4.05%) and liver (4.50% and 2.80% versus 15.5% and 2.05%; Figure 4C–D). No significant difference was detected between groups, though mice receiving EP-HSPCs trended higher. While we label these cells as EP-, that also means the cells were not exposed to the RNP complex nor to AAV6 transduction so any differences between EP-, EP+, and automation runs should be considered holistically and not simply as a consequence of EP.

To assess HSPC persistence in the BM and spleen, the two tissues to which these cells migrate in this model, we calculated CD34^+^ cells as a fraction of hCD45^+^ cells. In the BM, 7.46% of human cells were CD34^+^ following transplantation of edited HSPCs, versus 7.27% for unedited HSPCs. In the spleen, 3.50% of human cells were CD34^+^ in the edited HSPCs compared to 1.68% in unedited HSPCs (Figure 4E, Figure S2B). Neither difference in CD34^+^ populations was significant.

To assess differentiation into hematopoietic lineages, we tracked T cell, B cell, and monocyte populations in PB, BM, and spleen. B cells predominated across all tissues and groups, representing on average 71.0% of human cells in edited groups versus 70.4% in unedited groups. T cells represented 0.54% of human cells in edited groups versus 1.49% in unedited groups, and monocytes 2.22% versus 1.32% (Figure 4F). No significant difference in lineage distribution was detected between groups.

Finally, to determine whether editing persisted *in vivo*, we isolated genomic DNA from BM samples and sequenced it; because the primer sets were specific to the human genome, allelic distribution could be quantified against human alleles. For the automation runs, *CCR5* editing averaged 23.2% and 24.2% KI, and 59.9% and 73.2% KO (Figure 4G). These levels were comparable to those measured in geHSPCs before transplantation (21.1% and 24.7% KI, and 54.2% and 58.6% KO). Hence, at 12 weeks post-transplantation, geHSPCs from the final drug product remained detectable, showed no evidence of impaired engraftment *in vivo*, and confirmed that clinical scale manufacturing of geHSPCs in the shuttle retained the key critical quality attributes and biologic engraftment properties necessary for making a clinical grade product that would be infused into a patient.

## DISCUSSION

This study demonstrates that clinical-scale doses of geHSPCs can be produced on a fully automated, high-throughput manufacturing platform while meeting release criteria. Two independent runs starting from mPB CD34^+^ cells each yielded clinically relevant doses of the final drug product. High editing levels were preserved through process transfer to the automated platform and cell culture scale-up, and edited CD34^+^ cells retained the capacity to engraft in a humanized mouse model post-manufacturing. In showing that targeted gene integration can be ported onto a 5-day automated process starting with cryopreserved HSPCs to produce geHSPCs carrying multi-layered resistance to HIV, this work establishes the feasibility of automating a workflow that has, to date, depended on manual manufacturing that is costly and difficult to scale.

For a clinical autologous HSPCT program targeting HIV, both *CCR5* KO and KI efficiency are consequential. Because CCR5 serves as the co-receptor for R5-tropic HIV-1, cells that remain unedited constitute a reservoir of susceptible targets, and higher total KO levels are therefore expected to translate to better clinical outcomes.^21^ The *in vivo* total KO levels achieved here (60–73%) approached the thresholds (69–96% in humanized mice, 76–94% in non-human primates) predicted for HIV remission with *CCR5* editing alone, and exceeded those reported for *CCR5*-edited autologous products that have entered clinical trials, whether T cells (8.8%) or HSPCs (18–30%).^22–25^ Beyond *CCR5* KO, expression of a full-length bnAb in the B cell progeny that arise from geHSPCs adds a distinct mechanism of protection, neutralizing a broader set of strains including X4-tropic HIV-1, which do not depend on CCR5 for entry.^26^ HSPCs edited at the immunoglobulin locus reach clinically relevant bnAb levels at very low input thresholds (10–7,000 geHSPCs per mouse), well below the approximately 52,000–61,000 geHSPCs transplanted per mouse in this study.^3,4^ HSPCs with targeted antibody KI have yet to enter the clinic, so whether the *in vivo* KI levels observed at the *CCR5* locus (23–24%) will suffice to achieve remission remains to be determined. No HDR enhancers were used here, and their incorporation into future processes could further raise KI levels.^27–30^

These outcomes underscore the potential of geHSPCs as a one-time treatment for HIV and other chronic conditions. If geHSPCs engraft long-term, they could provide a lifelong supply of therapeutic antibody-secreting B cells. Recent reports have raised concerns that HDR impairs HSPC engraftment, though these studies used different repair templates (ssODN rather than AAV6) or higher AAV6 doses (10,000 versus 1,250 MOI).^31,32^ Transient inhibition of the p53 pathway restored engraftment, implicating downstream DNA damage responses such as inflammation and altered cell cycling, rather than editing itself, in the engraftment defect. Further, HSPCs were cultured at oxygen concentrations (20%) above those in the BM niche (3% O_2_), whereas HSPCs on the automated platform were cultured at oxygen concentrations similar to physiological conditions (5% O_2_).^33^ Although we detected a p53/p21 signature in geHSPCs three days post-manufacturing, a transient increase shown to resolve over time, these cells retained colony-forming capacity comparable to EP controls and maintained stable KI levels at 12 weeks post-transplantation, indicating that our process does not exceed the threshold of p53 activation associated with impaired engraftment.^34–36^ Automated handling of HSPCs, combined with lower viral vector doses and physiological culture conditions, may reduce cellular stress sufficiently to preserve the HSPC phenotype after editing. Clinical protocols will nonetheless warrant particular attention to ensure engraftment remains stable and undelayed following reinfusion of geHSPCs, for example by shortening the 5-day process or by delaying G-CSF treatment post-transplant.^36^ It is also possible that further refinements in the culture conditions such as described by Naseem et al might further increase HDR and increase the engraftability of geHSPCs.^37^

With over 40 million people living with HIV globally, automation will be essential to meeting demands for common conditions with high incidence. One Cell Shuttle could produce up to 778 geHSPC products a year using the process described here. An automated, high-throughput platform offers potential benefits beyond scale: reduced process variability, lower contamination risk, and a more consistent drug supply for patients awaiting transplantation. Together, these outcomes could reduce fixed costs, achieved by distributing personnel and facility across a larger number of batches, and variable costs, through reduced materials usage, decreased hands-on personnel time, and greater equipment utilization per project (Figure 5). These projections rest on assumptions about throughput and utilization that will be established over numerous runs, both for individual therapies in development and across therapies sharing the same process.

**Figure 5.**
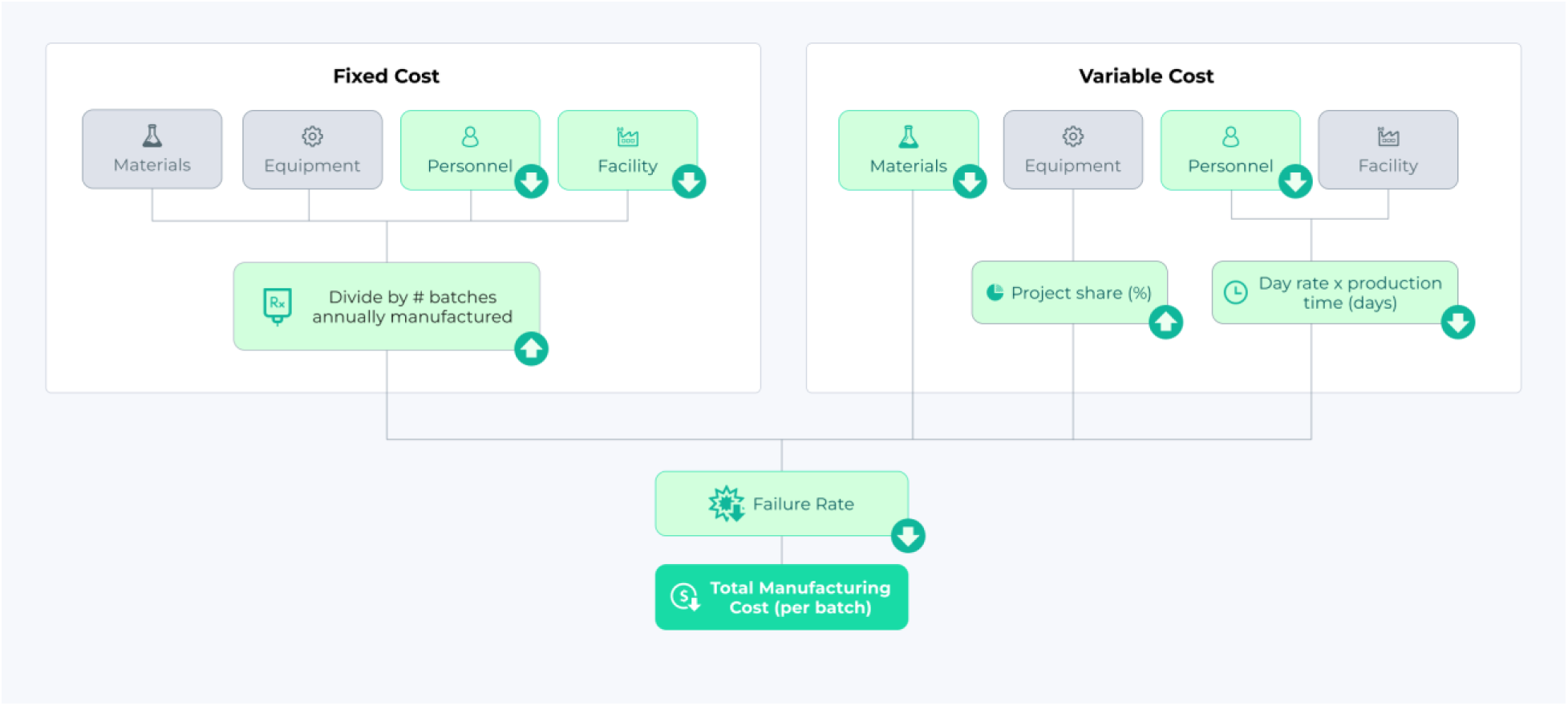
Cost model for geHSPC manufacturing on the automated platform. Total manufacturing cost per batch projected as the sum of fixed and variable cost components across four categories: materials, equipment, personnel, and facility. Fixed costs were allocated per batch by dividing annual costs by the number of batches manufactured annually. Variable costs were assigned by project share (%) for materials and equipment, and by day rate multiplied by production time (days) for personnel and facility. The combined cost was adjusted for failure rate to yield the total manufacturing cost per batch. Green shading and arrows denote categories in which the automated process is projected to reduce costs relative to conventional manufacturing.

Because *CCR5* functions as a safe harbor locus for transgene KI, the same editing approach could be repurposed across indications by only changing the therapeutic transgene.^38^ This *CCR5* safe harbor strategy is already being pursued for mucopolysaccharidosis type I and Gaucher disease.^39–41^ A platform approach in which the manufacturing process and sgRNA sequence remain fixed while the repair template varies could shorten development timelines and reduce cost by decreasing pre-clinical and chemistry, manufacturing, and controls (CMC) requirements, expanding access even for rare diseases with small patient populations.^42^ The Food and Drug Administration (FDA) has put forth guidance on a plausible mechanism framework that could open new approval pathways, though this has yet to become official policy.^43,44^ The Cell Shuttle has received Advanced Manufacturing Technology (AMT) designation from the FDA, reducing the regulatory burden for manufacturing methods, and a Platform Technology Designation could allow an initial product to establish criteria that support approval of subsequent products in future indications. By advancing *CCR5* safe harbor editing toward the clinic, this work could help lay the foundation for a broader set of HSPC gene therapies.

In summary, this study establishes the feasibility of producing clinical-scale doses of geHSPCs on an automated, high-throughput platform. By reducing the manual intervention and infrastructure that constrain conventional cell therapy manufacturing, this approach offers a route to expanding patient access to curative treatments worldwide.

### Limitations of the study

Several limitations constrain the conclusions of this work. Clinical-scale manufacturing was demonstrated in two automated runs using mPB CD34^+^ HSPCs from healthy donors. This number is sufficient to establish feasibility but not to characterize process variability. Run-to-run consistency in yield, editing, and product phenotype will require a larger series of runs to define. CD34^+^ isolation was performed upstream of the workflow described here; integration of isolation onto the automated platform, which has been validated for T cell workflows, is under development for HSPCs. CD34^+^ HSPCs from healthy donors may also behave differently from those mobilized from patients with HIV or other target indications, whose cell quality and yield can vary.

The *in vivo* studies carry the constraints of the model. Engraftment was assessed in NBSGW mice, in which hCD45^+^ reconstitution in PB was modest, consistent with prior reports for this strain.^19,20^ The xenotransplant model does not recapitulate human hematopoiesis, immune development, or the durability of engraftment over a clinically relevant timescale, and the 12-week endpoint does not address long-term repopulation by true long-term HSPCs.

The p53 pathway analysis was based on bulk RNA sequencing, which reports population-level averages and does not resolve whether the transcriptional response was uniform or confined to a subset of cells. The magnitude of p21 upregulation did not reach statistical significance with the number of runs performed, so the p53 signature is described but not quantified with confidence, and its functional consequence for engraftment was not directly tested.

Finally, this study measured editing frequencies, not antiviral function. The editing materials (sgRNA, AAV6) used in this process have previously been validated, and bnAb produced from edited B cells resulted in anti-HIV activity.^5,45^ Whether the *CCR5* KO and KI levels achieved translate to protection from HIV infection or to clinically meaningful bnAb secretion was not evaluated here though potential efficacy is supported by the recent Hartweger et al study demonstrating that <10% geHSPCs could generate inhibitory levels of bnAbs.^4^ The thresholds required for remission in humans remain undefined though the number of bnAbs and trough levels sufficient to maintain undetectable levels of HIV without antiretroviral therapy are beginning to be understood through trials using intermittent infusions of bnAbs into patients.^17,46^ In this approach, a one-time therapy might be sufficient to generate levels of bnAbs that are continuously maintained above suppressive levels without the risks of peaks and troughs.

## METHODS

### Key resources table

| REAGENT or RESOURCE | SOURCE | IDENTIFIER |
| --- | --- | --- |
| **Chemicals, peptides, and recombinant proteins** |  |  |
| UM171 | APExBIO | A89505 |
| SpyFi Cas9 10mg/mL | ALDEVRON | 9214-0.5MG |
| Recombinant Human SCF (Stem Cell Factor) | PreproTech | 300071MG |
| Recombinant Human TPO (thrombopoietin) | PreproTech | 300181MG |
| Recombinant Human FLT3-ligand | PreproTech | 300191MG |
| Recombinant Human IL-6 (Interleukin 6) | PreproTech | 200061MG |
| **Recombinant DNA** |  |  |
| CCR5 HDR templates - AAV6 for 10-1074 Ab, Vector: rAAV.GOI1.wnf029-101074 | Andelyn Bio | Feist et al. 2025<sup>5</sup> |
| **Oligonucleotides** |  |  |
| CCR5-targeting sgRNA | IDT | Feist et al. 2025<sup>5</sup> |
| **Experimental models:**<br>**Organisms/strains** |  |  |
| Mouse: NBSGW | The Jackson Laboratory | RRID:IMSR_JAX:026622 |
| **Biological samples** |  |  |
| CD34<sup>+</sup> HSPCs from mPB | Fred Hutchinson | RO04112B; RO04071 |
| CD34<sup>+</sup> HSPCs from mPB | AllCells | 889031123 (Male);<br>809044359 (Male);<br>3165590 (Male); 3165596 (Male) |
| CD34<sup>+</sup> HSPCs from mPB | StemCells | 230271401C (Male) |
| **Antibodies** |  |  |
| mCD45.1 (clone A20 BV650) | Biolegend | 110735;<br>RRID:AB_11124743 |
| hCD45 (clone HI30 BV421) | Biolegend | 304032;<br>RRID:AB_2561357 |
| hCD3 (clone HIT3a PerCP/Cy5.5) | Biolegend | 300328;<br>RRID:AB_1575008 |
| hCD19 (clone HIB19 PE-Cy7) | Biolegend | 982410;<br>RRID:AB_2715770 |
| hCD33 (clone P67.6 FITC) | Biolegend | 366620;<br>RRID:AB_2566422 |
| hCD34 (clone 581 APC) | Biolegend | 343510;<br>RRID:AB_1877153 |
| hCD90 (clone 5E10 PE) | Biolegend | 328110;<br>RRID:AB_893433 |
| hCD45RA (clone HI100 BV785) | Biolegend | 304140;<br>RRID:AB_2563816 |
| **Software and algorithms** |  |  |
| FlowJo v10.10.1 | TreeStar | RRID:SCR_008520 |
| Prism v11.0.0 | GraphPad | RRID:SCR_002798 |
| ICE 4.2 | EditCo | RRID:SCR_024508 |
| QuantaSoft | Bio-Rad | RRID:SCR_025321 |
| BioRender | BioRender | RRID:SCR_018361 |
| **Other** |  |  |
| CellGenix GMP Serum-free Stem Cell Growth Medium (SCGM) | CellGenix | 20802-0500 |
| MethoCult H4434 Classic | StemCell | 04444 |
| FcR Blocking Reagent, mouse | Miltenyi Biotec | 130-092-575 |
| FcR Blocking Reagent, human | Miltenyi Biotec | 130-059-901 |

### Ethics statement and HSPC source

Cryopreserved CD34^+^ hematopoietic stem cells (HSPCs) derived from mPB were purchased from StemCells, AllCells and Fred Hutchinson Cancer Center (Cooperative Center for Excellence in Hematology). All animal studies were conducted under protocols approved by the Stanford Institutional Review Board Committee and Administrative Panel on Laboratory Animal Care.

### *In vitro* culture and genome editing of CD34^+^ HSPCs

Cryopreserved CD34⁺ HSPCs were thawed into GMP-grade serum-free Stem Cell Growth Medium (SCGM, CellGenix) supplemented with 20 U/ml of penicillin, 20 mg/ml of streptomycin (Cytiva), 35 nM UM171 (APExBIO), and recombinant human (rh) Flt3 ligand (Flt3-L), interleukin-6 (IL-6), stem cell factor (SCF), and thrombopoietin (TPO) at 100 ng mL^-1^ each (PeproTech). Cells were cultured for 72 h at 37 °C in the Cell Shuttle prior to genome editing. Ribonucleoprotein (RNP) complexes were formed by incubating SpyFi Cas9 protein (Aldevron) with a CCR5-targeting sgRNA (IDT, sequence: 5’-CCUUCUUACUGUCCCCUUCU-3’) at a molar ratio of 1:2.7 for 15 min at room temperature. Cells were subsequently electroporated using the Cellares Electroporation Flowcell (Cellares) with parameters developed in-house, using Cellares’ proprietary EP buffer. Immediately following EP, cells were transduced with an AAV6 vector carrying a 10-1074 antibody-encoding donor template for homology-directed repair (HDR) at an MOI of 1,250 (rAAV.GOI1.wnf029-101074; Andelyn Biosciences). Cells were subsequently cultured at 37 °C for 48 h before harvest and formulation. Cell concentrations and viability were assessed using a Cellaca MX automated cell counter (Revvity).

### Commercial EP of CD34^+^ HSPCs

For the research-scale reference condition, 1×10⁶ CD34⁺ HSPCs were electroporated in a 100µL reaction using the 4D-Nucleofector System (Lonza) and P3 Primary Cell 4D-Nucleofector Kit. Cells were combined with precomplexed Cas9-sgRNA RNP immediately before EP and pulsed using program DZ-100. Following EP, cells were recovered in prewarmed HSPC culture medium and transferred to culture vessels containing the AAV.

### EP optimization in the Cellares Electroporation System

EP optimization was performed using human CD34^+^ HSPCs using Cellares pulser and Cellares proprietary EP buffer in a 2 mm gap EP cuvette (Thermo Fisher). Four EP conditions were evaluated: EP1 (400 V, 900 µs), EP2 (425 V, 700 µs), EP3 (425 V, 600 µs), and EP4 (450 V, 400 µs), each consisting of two pulses separated by a 4s interval. The optimized EP4 condition was subsequently evaluated across four independent healthy donors to assess donor-to-donor reproducibility.

### EP and cell culture scale-up

Cell culture and EP conditions were scaled-up across six stages, progressing from small-scale research conditions to fully automated clinical-scale production. At the research baseline (1x), cells were cultured in G-Rex (Wilson Wolf Manufacturing) 24-well vessels at 0.5×10^6^ cells/cm² (0.25×10^6^ cells/mL) in 4 mL of medium and electroporated in cuvette at a 100-µL scale. Following technology transfer and qualification (1x), cells were maintained in the same vessel format at an increased density of 2.5×10^6^ cells/cm^2^ (1.25×10^6^ cells/mL), with the 100-µL EP scale maintained. Vessel scale-up (5x) transitioned culture to G-Rex 6 vessels at 2.5×10^6^ cells/cm² (1.25×10^6^ cells/mL) in 200 mL of medium, with EP scaled to 0.5 mL (one batch Off-Cartridge in the Cellares flowcell). Platform translation (50x) moved culture into the Cellares Consumable Cartridge (CC), with cells seeded at 5×10^6^ cells/cm² (1.5×10^6^ cells/mL) in 165 mL of media and increased the EP scale to 5 mL across two batches in the Cellares flowcell. Clinical-scale benchtop production (125x) increased culture volume to 500 mL at a seeding density of 6.2×10^6^ cells/cm^2^ (1.25×10^6^ cells/mL) in the CC followed by further scale-up of EP to 12.5 mL across five batches, and clinical-scale automated production (125x) applied identical run parameters as a fully closed, automated process on the Cellares Cell Shuttle. Across scale-up, cells were resuspended prior to EP at a concentration of 1×10^7^ cells/mL for the research baseline or 5×10^7^ cells/mL for all subsequent stages.

### Cell Shuttle Automated Manufacturing

Cryopreserved CD34^+^ HSPCs were thawed, washed, and manually loaded into a Cellares Consumable Cartridge (CC) on day-3. The loaded cartridge was transferred to the Cellares Cell Shuttle where all subsequent processing was carried out over 5 days in a fully automated, functionally closed system with no human intervention with the exception of RNP and AAV6 formulation, which were prepared manually prior to loading. This included pre-EP culture, pre-EP wash, EP, media exchange, AAV6 transduction (TD), TD wash, culture, and final formulation. All reagent and media transfers throughout the automated manufacturing process were performed via sterile liquid transfer devices (SLTD).

For cryopreservation, cells were formulated in a 1:1 (v/v) mixture of Plasma-Lyte containing 2% human serum albumin (HSA) and CryoStor® CS10 and subjected to controlled-rate freezing.

### Quantification of edited alleles

Genomic DNA was extracted from HSPCs one day post-thaw using QuickExtract DNA Extraction Solution (LCG Biosearch Technologies). KO efficiency was quantified by indel analysis on Sanger sequencing results using inference of CRISPR edits (ICE 4.2; EditCo) as previously described. KI efficiency was quantified by the ratio of KI alleles to a reference gene using digital droplet PCR (ddPCR; Bio-Rad) as previously described.^5^ KO efficiency was normalized to account for KI alleles: % KO alleles = % indel / (100% – % KI alleles).

### Colony forming unit assays

One-day post-thaw, for each sample, 500 HSPCs were plated into two wells of a 6-well plate containing methylcellulose medium with cytokines (MethoCult H4434 Classic; StemCell Technologies). After 14 days in a hypoxic incubator (5% CO₂, 5% O₂) at 37°C, the wells were imaged using the STEMvision Hematopoietic Colony Counter (StemCell Technologies). Colonies were counted and scored with manual correction to determine the number of BFU-E, CFU-GM, and CFU-GEMM colonies.

### RNA-seq

RNA was extracted from HSPCs immediately post-thaw using Zymo DNA/RNA Shield (Sigma Aldrich). Bulk RNA was submitted for library preparation and sequencing (Plasmidsaurus). Deduplicated reads were mapped to genes to obtain counts, from which gene-level differential expression analysis was performed. Control (EP-and EP+) and edited (automation runs) results from the same run (>2.6 million uniquely mapped reads per sample) were grouped for comparison.

### Mouse engraftment studies

Adult (6–8-week-old) NOD.Cg-*Kit^W-41J^ Tyr*^+^ *Prkdc^scid^ Il2rg^tm1Wjl^*/ThomJ (NBSGW; Jackson Laboratories) mice were conditioned using a 10 mg/kg dose of busulfan delivered by intraperitoneal (IP) injection. 48 hours later, HSPCs were injected via intravenous injection (IV) into the tail vein. The condition of the mice was monitored for 8 weeks, after which biweekly (weeks 8, 10, and 12) PB was collected via retro-orbital bleeds into K2/EDTA tubes. At week 12, tissues were collected during necropsy. Both femurs and tibias were harvested for bone marrow extraction. Spleens and livers were processed into single-cell suspensions. Samples were hemolyzed using CTS ACK lysis buffer and split in half for genomic DNA collection and flow cytometry analysis.

### Flow cytometry

Samples for flow cytometry analysis were first stained for 20 minutes using Zombie-NIR Live/Dead Stain (BioLegend) at a 1:250 dilution. Samples were washed with FACS buffer containing 1 mM EDTA and 0.5% BSA. HSPCs were stained using the following antibodies: APC-anti-CD34, PE-anti-CD90, BV785-anti-CD45RA (Biolegend). For mouse samples, FcR blocking reagent for human and mouse (Miltenyi Biotec) was added for 10 minutes at a 1:25 dilution, followed by BV650-anti-mouse-CD45.1, BV421-anti-CD45, PerCP/Cy5.5-anti-CD3, PE-Cy7-anti-CD19, and FITC-anti-CD33 (Biolegend). All antibodies were incubated for 30 minutes at 1:50 dilutions, then washed with FACS buffer and run on a FACSAria II or an LSRFortessa (BD Biosciences). Results were gated on live, single cells and analyzed using FlowJo (TreeStar, v10.10.1).

### Statistical analysis

All statistical analysis was performed using Prism v11.0.0 software (GraphPad).

### Data availability

Data are available from the corresponding author upon request.

## SUPPLEMENTAL INFORMATION

Document S1. Figures S1-S2. Table S1.

## ACKNOWLEDGEMENTS

The authors thank Dr. Andrea Leonardi for supporting figure design, Dr. Yangzi Isabel Tian for reviewing the manuscript, members of the Porteus lab for assisting with mouse necropsies, and Stanford Institute for Stem Cell Biology and Regenerative Medicine FACS Core for providing access to flow cytometers. This work was supported by the Stanford Innovative Medicines Accelerator (IMA-1386) and in part by Cooperative Centers of Excellence in Hematology (NIDDK Grant #DK106829). N.A. was supported by the Propel Postdoctoral Scholars Program at Stanford University School of Medicine, the American Society of Hematology (ASH), HIP Fellow Award, and by the Learners to Leaders in Urology, Nephrology, and Non-Cancer Hematology Program (LAUNCH; NIDDK Grant #DK139565). S.E.L. was supported by the Stanford Medical Scientist Training Program and NIH F30 Individual Predoctoral Fellowship (NHLBI #HL178496). F.K.E. was supported by the Stanford Knight-Hennessy Scholarship, the Paul & Daisy Soros Fellowship for New Americans, and the Hertz Fellowship. M.H.P. was supported by the Sutardja Chuk Professorship in Definitive and Curative Medicine and the Laurie Kraus Lacob Translational Medicine endowment.

## AUTHOR CONTRIBUTIONS

J.M.P.C., N.A., S.E.L., F.U., and M.H.P. conceptualized the project and designed experiments. D.M., K.S., V.R., A.B., A.S., J.F., Z.L., K.R., and J.F. performed and collected data from process development and automation runs. J.M.P.C., N.A., and F.K.E. performed and collected data from mouse engraftment studies. J.M.P.C. analyzed data and drafted the manuscript. N.A., S.E.L., F.U., and M.H.P. reviewed and revised the manuscript. All authors have read and agreed to the current version of the manuscript.

