## Supplemental information for "End-To-End, Fully Automated Manufacturing of Hematopoietic Stem and Progenitor Cell Therapy Via*CCR5*-Targeted Integration"

### SUPPLEMENTAL FIGURES

#### Figure S1.

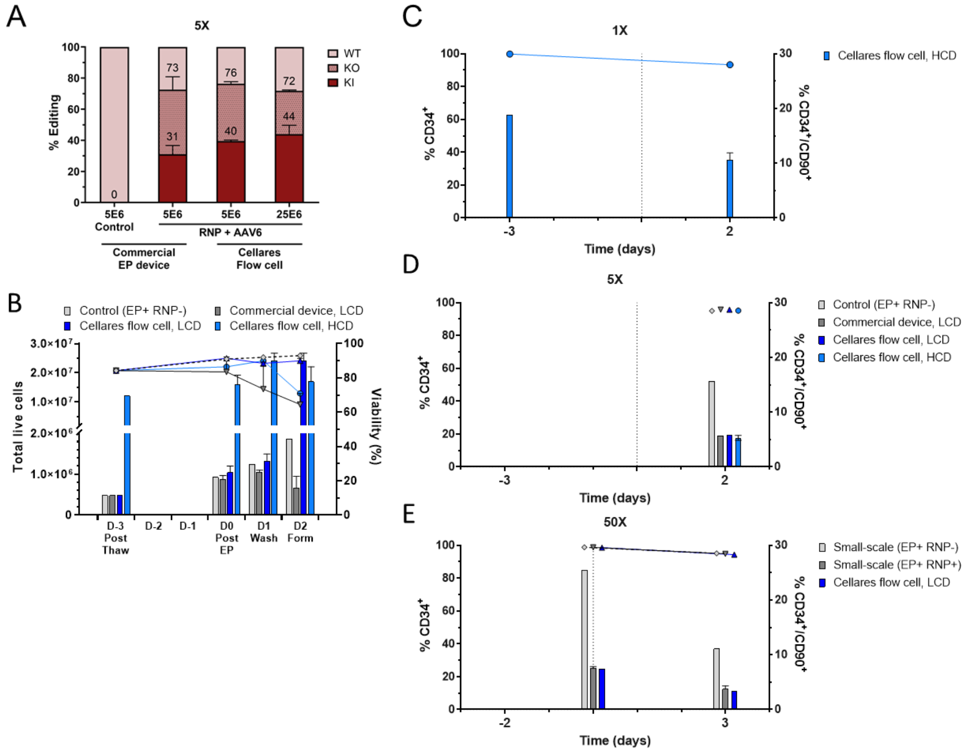

**Figure S1. Intermediate EP scale and HSPC purity through scale-up.**

(A, B) Editing performance and viability at intermediate EP scale (5x: 500 µL) at HCD ($5x10^7$ cells/mL). (A) KO and KI efficiencies. (B) Total live cells and viability across process time points. (C-E) Frequency of $CD34^+/CD90^+$ (bars, left axis) and $CD34^+$ (lines/points, right axis) populations at different time points. (C) HSPC purity at 1x scale. (D) HSPC purity at 5x scale. (E) HSPC purity at 50x scale. Mean and SD are plotted; n = 1-2 replicates.

**Figure S2.**

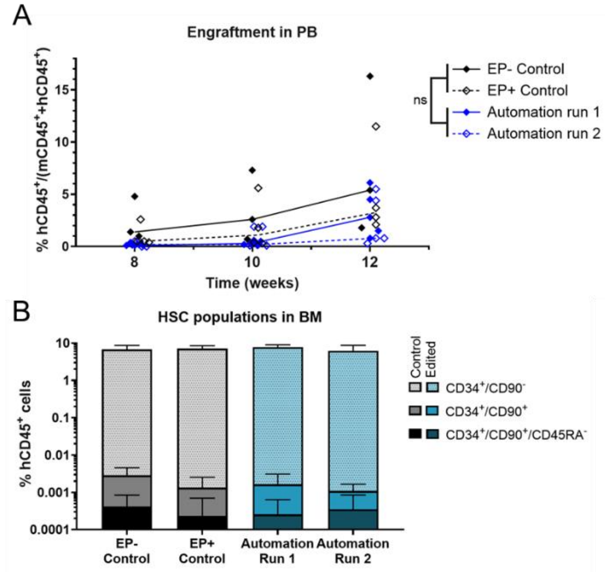
A
Engraftment in PB
% hCD45+/(mCD45+ +hCD45+)
Time (weeks)
ns
EP- Control
EP+ Control
Automation run 1
Automation run 2
B
HSC populations in BM
% hCD45+ cells
Control
Edited
CD34+/CD90-
CD34+/CD90+
CD34+/CD90+/CD45RA-
EP- Control
EP+ Control
Automation Run 1
Automation Run 2

**Figure S2. Peripheral blood engraftment and HSC populations in bone marrow of NSBGW mice.**

(A) Engraftment in PB obtained by retro-orbital bleeds between weeks 8 and 12 as measured by flow cytometry on human and mouse $CD45^+$ cells. (B) Frequency of $CD34^+/CD90^-$, $CD34^+/CD90^+$, and $CD34^+/CD90^+/CD45RA^-$ populations in BM at necropsy. Mean and SD are plotted; n = 3-5 mice; significance calculated by unpaired two-tailed t test at each time point.
